# An IL-21R STAT hypomorph circumvents functional redundancy in germinal center responses

**DOI:** 10.1101/2025.11.14.688418

**Authors:** Christoph Jandl, Joanna Warren, Samantha Owens, Marcel Batten, Howard Wang, Cecile King

## Abstract

Interleukin-21 receptor engagement initiates a Janus Kinase (JAK) - signal transducer and activator of transcription (STAT) signalling cascade that activates several STAT proteins that drive the germinal center response. However, the relative effects of IL-21 on individual STAT proteins during the differentiation of T follicular helper cells and germinal center B cells has been difficult to distinguish. Here, we characterise a novel mutation in the interleukin-21 receptor (IL-21R^EINS^) that creates a unique defect in the activation of STAT1. Our findings provide evidence that IL-21 mediated activation of STAT1 has a nonredundant role in the differentiation of T follicular (Tfh) cells following T dependent immunisation. IL-21R^EINS^ Tfh cells were even more impaired than Tfh cells genetically deficient in IL-21R, questioning our current understanding of the role of IL-21 derived from protein knockout mice. The observation that functional compensation fails in the presence of the IL-21R hypomorph provides insight into how underlying compensation can impact our interpretation of a complex biological system.

## Introduction

Signals delivered by cytokines are known to shape the humoral immune response to T-dependent antigen, acting on multiple cell subsets involved in this process (*1*). Interleukin-21 signals through the IL-21 receptor (IL-21R), which is a dimeric receptor consisting of the IL-21Rα chain and the γc belonging to the family of common γ chain (γc) receptors (*2–6*). IL-21:IL-21R engagement leads to the activation of Janus kinase (JAK)1 and JAK3, which, in turn, results in the phosphorylation of Signal Transducer and Activator of Transcription (STAT)1, STAT3 and, to a lesser extent, STAT5 (*4, 7–9*). Previous studies have shown the importance of signalling mediated by the cytokine interleukin (IL)-21 for the differentiation of T follicular helper (Tfh) cells and germinal centre (GC) B cells (*10–15*), with some studies questioning a T cell intrinsic role for IL-21 (*12, 13, 16*).

IL-21, produced in abundance by Tfh cells (*11*), activates STAT1 and STAT3 in both Tfh cells and B cells, influencing humoral immune responses. IL-21 co-stimulates the antigen signal in CD4^+^ T cells and can induce the production of IL-21, amplifying its own effects (*17*). IL-21 mediated activation of the JAK-STAT pathway is required for the establishment of long-lasting immune responses in both humans and mice (*7, 10, 11, 18, 19*). Analyses of IL-12–induced expression of IL-21 by human CD4^+^ T cells demonstrated a nonredundant role of *STAT3* in human Tfh cell differentiation, whereas *STAT1* was dispensable (*20*). By contrast, in mice, STAT1 activity was required for Bcl6 induction and early Tfh differentiation following LCMV infection (*21*). In B cells, STAT3 plays a critical role in generating memory cells and plasma cells (PCs) from naïve murine precursors *in vivo* and in human cells in response to IL-10 and IL-21 *in vitro* (*7*). Analyses of mice containing a selective *Stat3* deficiency in CD4^+^ T cells demonstrate reduced Tfh and GC B cell differentiation following viral infection and *Stat3*-deficient Tfh cells exhibited a transcriptome similar to Th1 cells (*22*).

Similar to IL-21, the cytokine IL-6 induces the activation of STAT1 and STAT3 and promotes the differentiation of Tfh cells. A reduction in the magnitude of the GC response and Tfh cell numbers can be observed in mice deficient in the cytokine IL-6 (*Il6^−/−^* mice) (*10, 23*), and the absence of both IL-6 and IL-21 *in vivo* led to an even greater decrease in Tfh cell numbers in response to infection that the absence of either cytokine alone, indicating that IL-21 and IL-6 have redundant functions for Tfh cell differentiation (*16, 24*). Both IL-6 and IL-21 drive the expression of molecules important for Tfh cell differentiation including the Tfh cell master transcription factor Bcl-6 (*21, 25–27*). Furthermore, IL-6 has not only been shown to compensate for a lack of IL-21 but is also able to induce the production of IL-21, further positively influencing Tfh cell development (*28*). In this context, the degree to which IL-21 is crucial for Tfh cell differentiation is affected by the infection (*16*) and the type of adjuvant applied to immunisation, where IL-6 and type-I interferon inducing Pathogen-Associated Molecular Patterns (PAMPS), such as found in sheep red blood cell (SRBC) immunisation (*29*), might compensate for the lack of IL-21 signalling in Tfh cells (*11, 14, 30*).

Previous studies analysing *STAT3* and *STAT1* mutations in humans and *Stat1*- and *Stat3*-deficient mice and have shown that both STAT1 and STAT3 are important for IL-21 mediated gene regulation in CD4^+^ T cells (*31*), with *Stat3* heterozygosity adequate to drive a profound effect on Th17 cell differentiation (*32*). However, the relative importance of the activation of STAT1 and STAT3 downstream of the IL-21R for the GC reaction remains incompletely understood. In this study, which is based in part upon the dissertation of Christoph Jandl (*33*), we analyse the effect of a single point mutation in the cytosolic region of the IL-21R that limits IL-21 mediated STAT1 activation and show that this partial loss of function mutation inhibits the differentiation of Tfh cells and reduces the magnitude of the GC reaction to T-dependent antigen. Tfh cells that harboured a defective IL-21R were more impaired in response to sheep red blood cell (SRBC) immunization than Tfh cells deficient in the IL-21R, offering insights into IL-6 mediated compensatory mechanisms that occur when the protein is completely ablated.

## Results

### The *einseitig* (*EINS*) mutation in IL-21R leads to a unique STAT1 signalling defect

Tfh cell differentiation in mice made genetically deficient in the full-length IL-21R protein has revealed that IL-6 can, to varying degrees, compensate for the absence of IL-21:IL-21R signalling (*16, 24*). We queried whether the same level of compensation would operate in the presence of a mutant IL-21R. To test this, we analysed a novel mouse line that that was generated on a C57BL/6 background by N-ethyl-N-nitrosourea (ENU) mutagenesis, a method to chemically induce point mutations in the DNA of murine spermatogonial stem cells (*34*). A single point mutation in IL-21R (Fig. 1A), that we named *einseitig,* was identified in these mice (*Il21r^EINS^* mice), leading to a change from the highly conserved Aspartic acid to Valine on amino acid position 375 of the receptor protein (Fig. 1B). This change from a negatively charged to a neutral amino acid in the cytoplasmic tail region that controls STAT activation resulted in a PolyPhen-2 score of 0.99, predicting a high likelihood of a functional disruption of the IL-21R.

**Figure 1.**
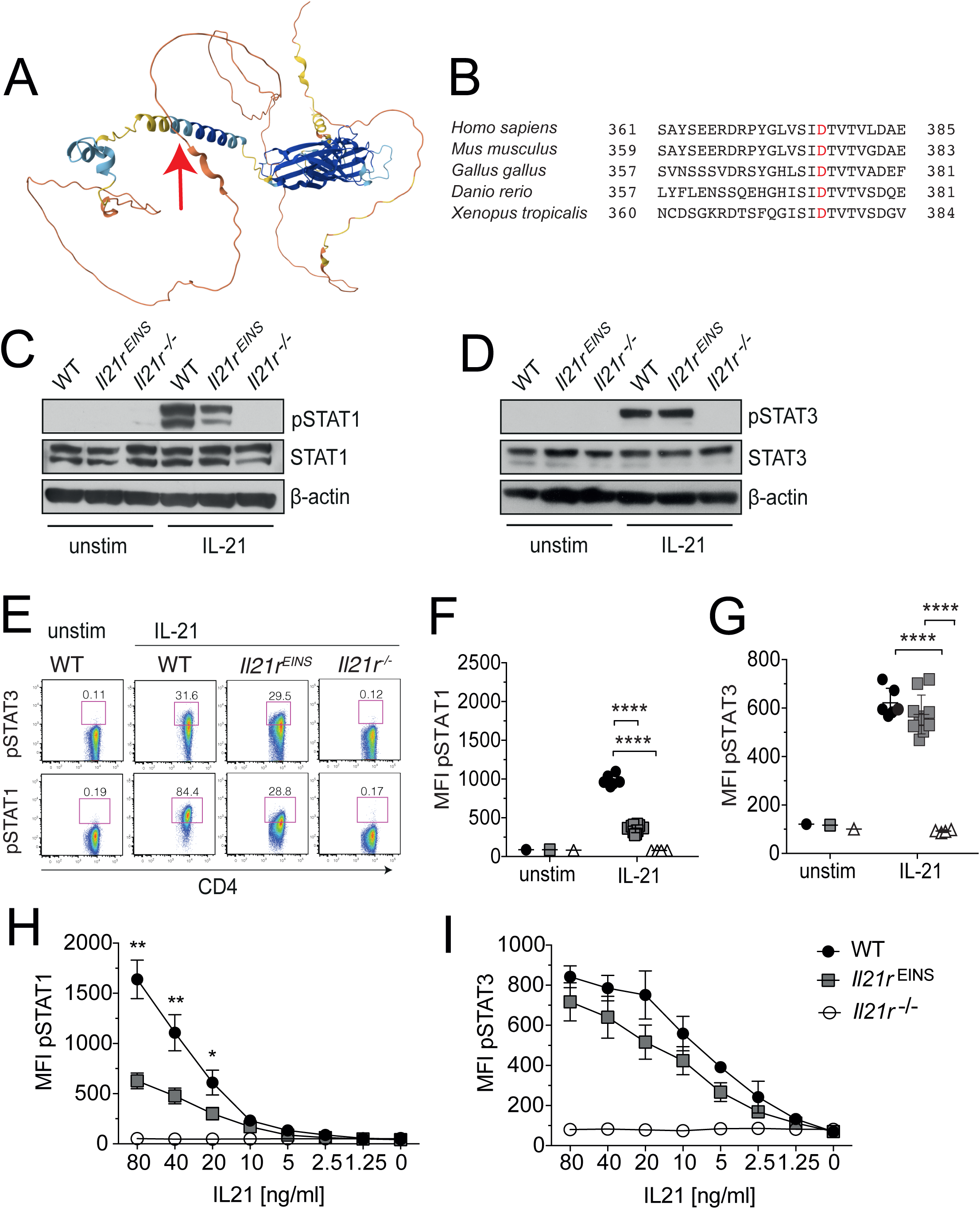
Selective defect in IL-21 induced STAT1 activation in *Il21r^EINS^* cells. The structure of murine IL-21R generated by alphaFOLD, showing the *einseitig* mutation leading to a change from Aspartic acid to Valine at amino acid position 375 (A). The Aspartic acid at amino acid position 375 is highly conserved (B). Splenocytes from naïve WT, *Il21r^EINS^* and *Il21r−/−* mice were stimulated with 80ng/ml rmIL-21, rmIL-6 or incubated in media for unstimulated controls for 15min. Western blots of total splenocytes either unstimulated or stimulated for 15 mins with rmIL-21, lysed with RIPA buffer and stained with antibodies against phosphorylated (p)STAT1 (PY701) and STAT1 (C), pSTAT3 (PY705) and STAT3 (D) are shown. For flow cytometric analysis we gated on TCRβ^+^ CD4^+^ cells (E) to assessed levels of intracellular P-STAT1 (F) and P-STAT3 (G) expression on CD4^+^ T cells after stimulation for 15 mins *in vitro* with recombinant murine IL-21. Titration curves showing phosphorylation of (H) STAT1 and (I) STAT3 determined by flow cytometry and FACS analyses after 15 min stimulation of spleen derived CD4^+^ T cells with rmIL-21. Graphs show individual mice, Data are shown as mean ± SD, representative of 4 similar individual experiments. Statistical significance was assessed by one-way ANOVA using Bonferroni’s multiple comparisons test; *\*p < 0.05; **p < 0.01; ***p < 0.001; ****p < 0.0001*.

Since the engagement of IL-21R with its ligand IL-21 predominantly leads to phosphorylation of both STAT1 and STAT3 (*4, 35*), we first characterized the impact of the *einseitig* mutation (Fig. 1A) on the phosphorylation of STAT proteins in response to IL-21. Splenocytes from WT, *Il21r^EINS^* and *Il21r^−/−^* mice were stimulated with recombinant murine (rm) IL-21 for 15 minutes *in vitro* and the phosphorylation of STAT1 and STAT3 assessed by Western blot. IL-21 dependent STAT1 phosphorylation was reduced in *Il21r^EINS^* splenocytes compared with WT (Fig. 1C), whereas STAT3 phosphorylation in *Il21r^EINS^* splenocytes in response to IL-21 appeared intact (Fig. 1D). Control *Il21r^−/−^* splenocytes showed an absence of IL-21 dependent phosphorylation of both STAT1 (Fig. 1C) and STAT3 (Fig. 1D).

To further interrogate the STAT1 defect, splenocytes from WT, *Il21r^EINS^*and *Il21r^−/−^* mice were stimulated with rmIL-21 *in vitro* and the phosphorylation levels of STAT1 and STAT3 in CD4^+^ T cells were assessed by intracellular immunostaining and flow cytometry (Fig 1E). Consistent with our Western blot analyses, we observed a marked decrease in the mean fluorescence intensity (MFI) of IL-21 induced STAT1 phosphorylation in *Il21r^EINS^* CD4^+^ T cells compared with WT CD4^+^ T cells, whereas no pSTAT1 could be detected in *Il21r^−/−^* CD4^+^ T cells (Fig. 1E and Fig. 1F). By contrast, the levels of STAT3 phosphorylation were comparable in *Il21r^EINS^* and WT CD4^+^ T cells. As expected, no pSTAT3 could be detected in *Il21r^−/−^* cells in response to rmIL-21 (Fig. 1E and Fig 1G).

To exclude the possibility that the ability of the mutant IL-21R to mediate STAT1 and STAT3 phosphorylation was dependent on the intensity of the IL-21-mediated signal, we also assessed pSTAT1 and pSTAT3 levels in response to decreasing concentrations of rmIL-21. Corresponding to observations with a single concentration of rmIL-21, the levels of pSTAT1 in *Il21r^EINS^*CD4^+^ T cells were reduced in comparison to WT across a wide range of rmIL-21 concentrations (Fig. 1H), whereas the levels of phosphorylated STAT3 in CD4+ T cells were significantly less affected by the *Il21r^EINS^* mutation (Fig. 1I). As anticipated *Il21r^−/−^*CD4^+^ T cells did not activate either STAT1 or STAT3 following stimulation with rmIL-21 (Fig. 1H and Fig. 1I). Since IL-6 activates both STAT1 and STAT3 through a different receptor than IL-21, stimulation with IL-6 served as a positive control for STAT activation (*21, 24, 36*). Despite a STAT1 defect in response to IL-21, percentages of *Il21r^EINS^*CD4^+^ T cells showed equivalent STAT1 activation to WT cells when stimulated with rmIL-6 (Figure S1A). The percentages of CD4+ T cells that activated STAT3 in response to IL-6 were also similar between *Il21r^EINS^*, *Il21r^−/−^* and WT CD4^+^ T cells (Figure S1B).

In addition to STAT1 and STAT3, IL-21 has also been shown to activate STAT5, albeit to a lesser extent than IL-2 (*8*). To assess STAT5 phosphorylation in *Il21r^EINS^* CD4^+^ T cells, we stimulated splenocytes from WT, *Il21r^EINS^* and *Il21r^−/−^*mice either with rmIL-6, rmIL-21 or rmIL-2 *in vitro* and subsequently measured the expression levels of pSTAT5 in CD4^+^ T cells by flow cytometry. Expression levels and percentages of pSTAT5 in WT and *Il21r^EINS^*CD4^+^ T cells were equivalent in response to all three stimulatory conditions (Figure S1C and S1D). Furthermore, titration of rmIL-21 activated STAT5 to a similar extent in WT and *Il21r^EINS^* CD4^+^ T cells shown both as an increased intensity of pSTAT5 (Figure S1E) and percentage of CD4^+^ T cells containing pSTAT5 (Figure S1F), whereas no effect was observed in *Il21r^−/−^* cells. Taken together, our findings provide evidence that the *einseitig* mutation in IL-21R leads to a distinct STAT1 activation defect that does not affect either STAT3 or STAT5 activation in response to IL-2. *Il21r^EINS^*mice thus provide a unique opportunity to distinguish the contribution of IL-21 mediated STAT1 activation in the germinal centre response.

### The *einseitig* mutation leads to a defective GC response to T-dependent antigen

Prior to immunisation, lymphoid and myeloid cell populations in the thymus and the spleen of naïve *Il21r^EINS^* mice were analysed, indicating no major impact of the *einseitig* mutation on the development of lymphoid cell populations when compared with WT mice (Figure S2). To assess the influence of the *einseitig* mutation on the GC reaction, we compared the response of *Il21r^EINS^*, *Il21r^−/−^* and WT mice following immunisation with the T-dependent antigen, sheep red blood cells (SRBC). Histological analyses of spleen sections from mice 7 days after SRBC immunization showed that GCs were still able to form when IL-21 dependent STAT1-signaling was diminished (Fig. 2A). However, the reduction of IL21-mediated STAT1 phosphorylation led to decreased percentages (Fig. 2B and 2C) and numbers (Figure S3) of FoxP3^−^ CXCR5^high^ PD-1^high^ CD4^+^ Tfh cell population 7 days after SRBC immunization despite intact STAT3 phosphorylation downstream of the IL-21R. Interestingly, this defect was more pronounced in the presence of limited signalling through the mutant IL-21R, since *Il21r^−/−^* Tfh cells did not exhibit an equivalent reduction in response to SRBC immunization (Fig. 2C and Figure S3). These data suggest that whilst IL-6 has been shown to compensate for a lack of IL-21 in Tfh cell differentiation (*16, 24*), no such redundancy operated when a defective IL-21R was present.

**Figure 2.**
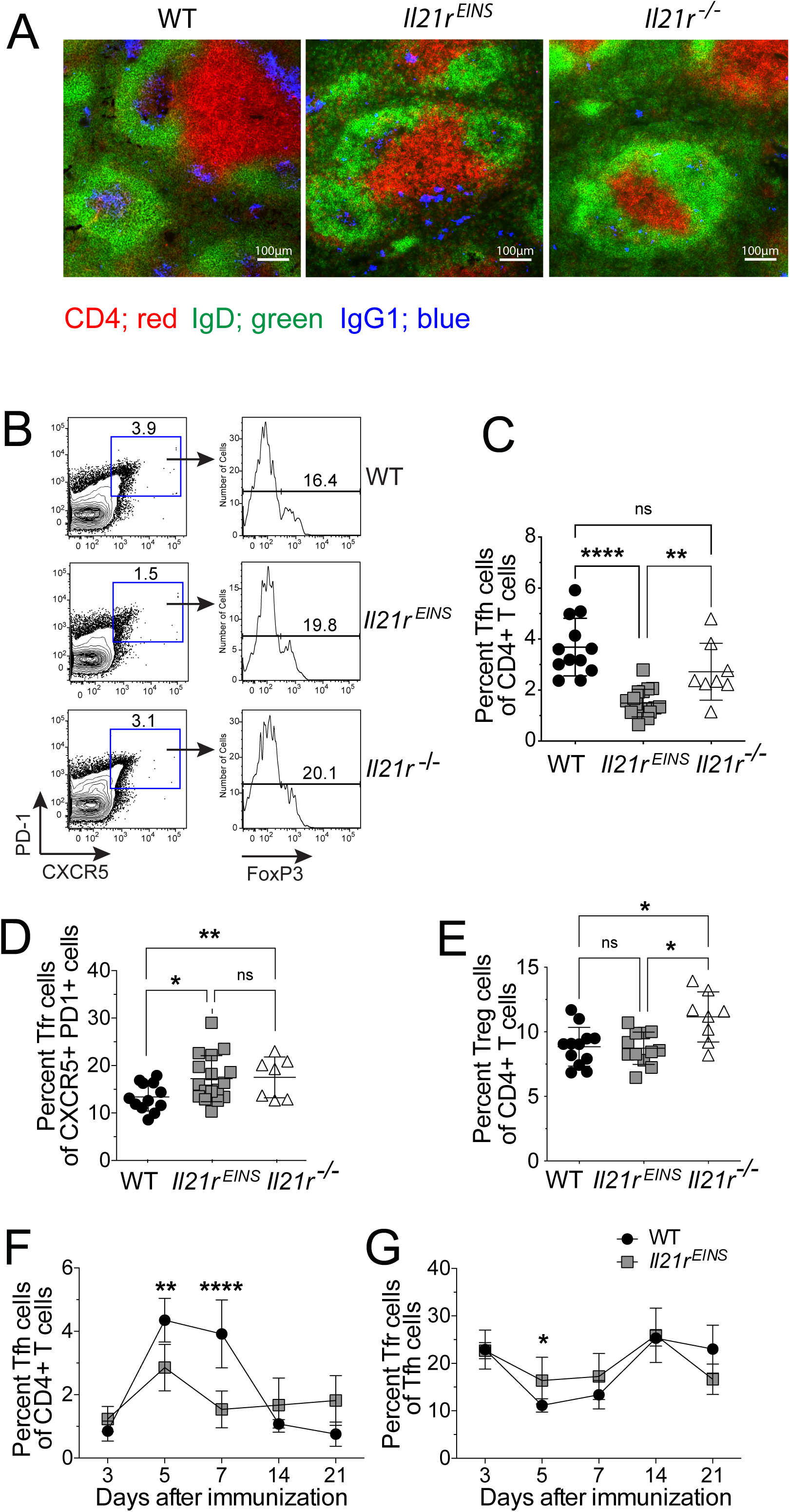
The *Il21r^EINS^* mutation reduces the differentiation of T follicular helper cells. WT, *Il21r^−/−^* and *Il21r^EINS^*mice were immunized with SRBC and the kinetics of the immune response was observed by analysing the response on the timepoints indicated. Analysis of the GC response in the spleen in WT, *Il21r^−/−^* and *Il21r^EINS^*mice on day 7 following SRBC immunization. (A) Histological sections of the spleen on day 7 of SRBC immunisation showing CD4+ cells (red), IgD (green) and IgG1 (blue). (B) FACS dot plots shows gating strategy for CXCR5^high^ and PD-1^high^ CD4^+^ T cells and FoxP3^+^ Tfr cells. (C) Percentage of T follicular helper (Tfh) cells (CXCR5^high^ PD-1^high^ FoxP3^−^ CD4^+^ T cells) as a percentage of CD4^+^ T cells population. (D) Percentage of FoxP3^+^ T follicular regulatory (Tfr) cells of the CXCR5^high^ PD-1^high^ CD4^+^ T cell population. (E) Percentage of FoxP3^+^ Treg cells within the CD4^+^ T cell population. Values shown from individual mice n=4-8 per group, from 2 pooled experiments including means +/- SD with 2 experimental replicates. Kinetics of the (F) T follicular helper (Tfh) cell response and (G) the T follicular regulatory (Tfr) cell response over a 21-day time-course data is shown as means +/- SD, n=4-8 with 2 experimental replicates. Statistical significance was assessed by one-way ANOVA using Bonferroni’s multiple comparisons test; *\*p < 0.05; **p < 0.01; ***p < 0.001; ****p < 0.0001*.

We have shown previously that the FoxP3^+^ CXCR5^high^ PD-1^high^ CD4^+^ Tfr cell population was increased in *Il21r^−/−^* mice (*37*). We similarly observed an increase in the percentages (Fig. 2D), but not the numbers (Figure S3), of Tfr cells in *Il21r^EINS^* mice in comparison to WT mice 7 days after SRBC immunization. However, FoxP3^+^ CD4^+^ T regulatory cells (Tregs) were observed at similar frequencies in wild-type and *Il21r^EINS^* mice, but were increased in *Il21r^−/−^*mice, following SRBC immunization (Fig. 2E). Assessment of the kinetics of T follicular cells showed the percentages of Tfh cells were significantly decreased 5 and 7 days after SRBC immunization (Fig 2F), and the percentage of Tfr cells showed a slight increase 5 days after SRBC immunization (Fig. 2G) in *Il21r^EINS^* mice compared with WT mice. Taken together, these findings suggested that IL-21 mediated STAT1 phosphorylation supported the differentiation of Tfh cells and inhibited the differentiation of Tfr cells.

Consistent with these findings, the percentages of *Il21r^EINS^*GC B cells (Fig. 3A) and IgG1^+^ FAS^+^ B cells (Fig. 3B) were significantly reduced relative to WT GC B cells but were slightly increased when compared with *Il21r^−/−^* GC B cells. These data were supported by a decrease in absolute numbers of GC B cells (Fig. 3C) and IgG1^+^ FAS^+^ B cells (Fig. 3D) in both *Il21r^EINS^*and *Il21r^−/−^*mice compared with WT mice. Further analyses of the kinetics of the GC populations by flow cytometry from day 3 to day 21 after SRBC immunisation showed that the percentages of GC B cells were decreased in *Il21r^EINS^* mice relative to their WT counterparts on day 5 and 7 (Fig. 3E).

**Figure 3.**
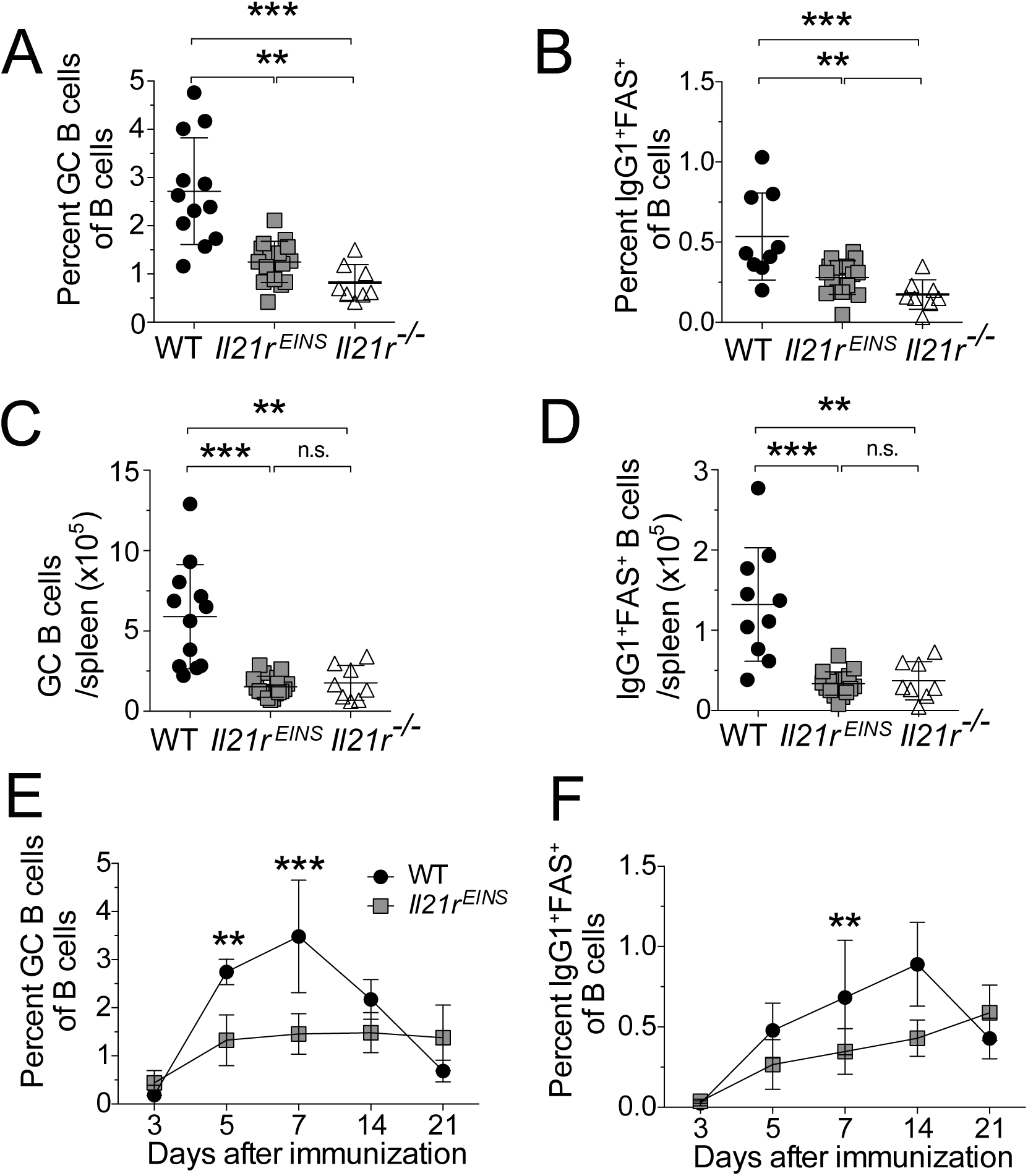
The *Il21r^EINS^* mutation reduces germinal centre output. WT, *Il21r^−/−^* and *Il21r^EINS^*mice were immunized with SRBC, and the kinetics of the immune response was observed by analysing the response on the timepoints indicated. FAS^+^ GL7^+^ GC B cells as a percentage of total B cells (A), IgG1^+^ FAS^+^ C cells as a percentage of total B cells (B). Absolute numbers of GC B cells (C) and IgG1^+^ FAS^+^ B cells (D) per spleen. Analysis of splenocytes on day 7 following SRBC immunization. Data are shown as mean +/- SD, n=5-8 mice with 2 experimental replicates. Kinetics of the GC B cell response (E) and the IgG1^+^ FAS^+^ B cell response (F) in the spleen of WT and *Il21r^EINS^* mice. Values are shown from individual mice, including means +/- SD from 3 experimental repeats where n=4-5 per group. Statistical significance was assessed by one-way ANOVA using Bonferroni’s multiple comparisons test; *\*p < 0.05; **p < 0.01; ***p < 0.001; ****p < 0.0001*.

Despite an increase in the IgG1^+^ FAS^+^ B cell population from day 3 in response to immunization, the percentages of *Il21r^EINS^* IgG1^+^ GC B cells were significantly lower relative to WT cells 7 days after SRBC immunisation (Fig. 3F). These findings demonstrate that IL-21 mediated STAT1 activation supports the differentiation of GC B cells following SRBC immunisation.

### IL-21 dependent activation of STAT1 has a T cell intrinsic component

To determine the cell intrinsic effects of the STAT1-specific IL21-signaling defect caused by the *einseitig* mutation on GC populations, we transferred equal amounts of bone marrow (BM) cells from WT mice expressing the congenic marker CD45.1 and BM from either *Il21r^EINS^*or WT expressing CD45.2 into lethally irradiated RAG-1 deficient mice. After 8 weeks of rest to allow for reconstitution, we checked for chimerism in the blood and subsequently immunized the BM chimera mice with SRBC and the relative contribution of WT and *Il21r^EINS^* cells to T and B cell populations was examined. Control chimeras reconstituted with 50% WT CD45.1 BM and 50% WT CD45.2 BM exhibited similar contributions of CD45.1^+^ and CD45.2^+^ cells to the total CD4^+^ T cell, Treg cell, Tfr cell, and Tfh cell populations after immunization with SRBC (Figure S4), therefore excluding a potential impact of the congenic markers on reconstitution.

Analyses of the spleens of chimeric mice following immunization with SRBC showed that CD45.2^+^ *Il21r^EINS^* cells contributed only an average of 41.2% of the total donor CD4^+^ T cell population, suggesting that the *Il21r^EINS^* mutation negatively affected CD4^+^ T cell proliferation or survival (Fig. 4A and Fig. 4B). However, the *Il21r^EINS^* Tfh cell population showed a small but consistent decrease relative to CD45.2^+^ CD4^+^ T cells deriving from *Il21r^EINS^* donor mice, indicating decreased differentiation of Tfh cells (Fig. 4A and Fig. 4B). By contrast, the *Il21r^EINS^*FoxP3^+^ Tfr population showed an increased proportion relative to CD4^+^ T cells deriving from *Il21r^EINS^* donor mice (Fig. 4A and Fig. 4C). Parallel analyses of the T-cell subsets within each donor population derived either from CD45.1^+^ WT or CD45.2^+^ *Il21r^EINS^* donor BM within individual chimeric mice confirmed a decrease of *Il21r^EINS^* Tfh cells (Fig. 4D) and showed small increases in both CD45.2^+^ *Il21r^EINS^*FoxP3^+^ Tfr cells (Fig. 4E) and Treg cells (Fig. 4F) in comparison with WT cells. Taken together, these findings provide evidence for a cell intrinsic contribution of IL-21 dependent STAT1-signaling to the differentiation of both Tfh cells and Tfr cells.

**Figure 4.**
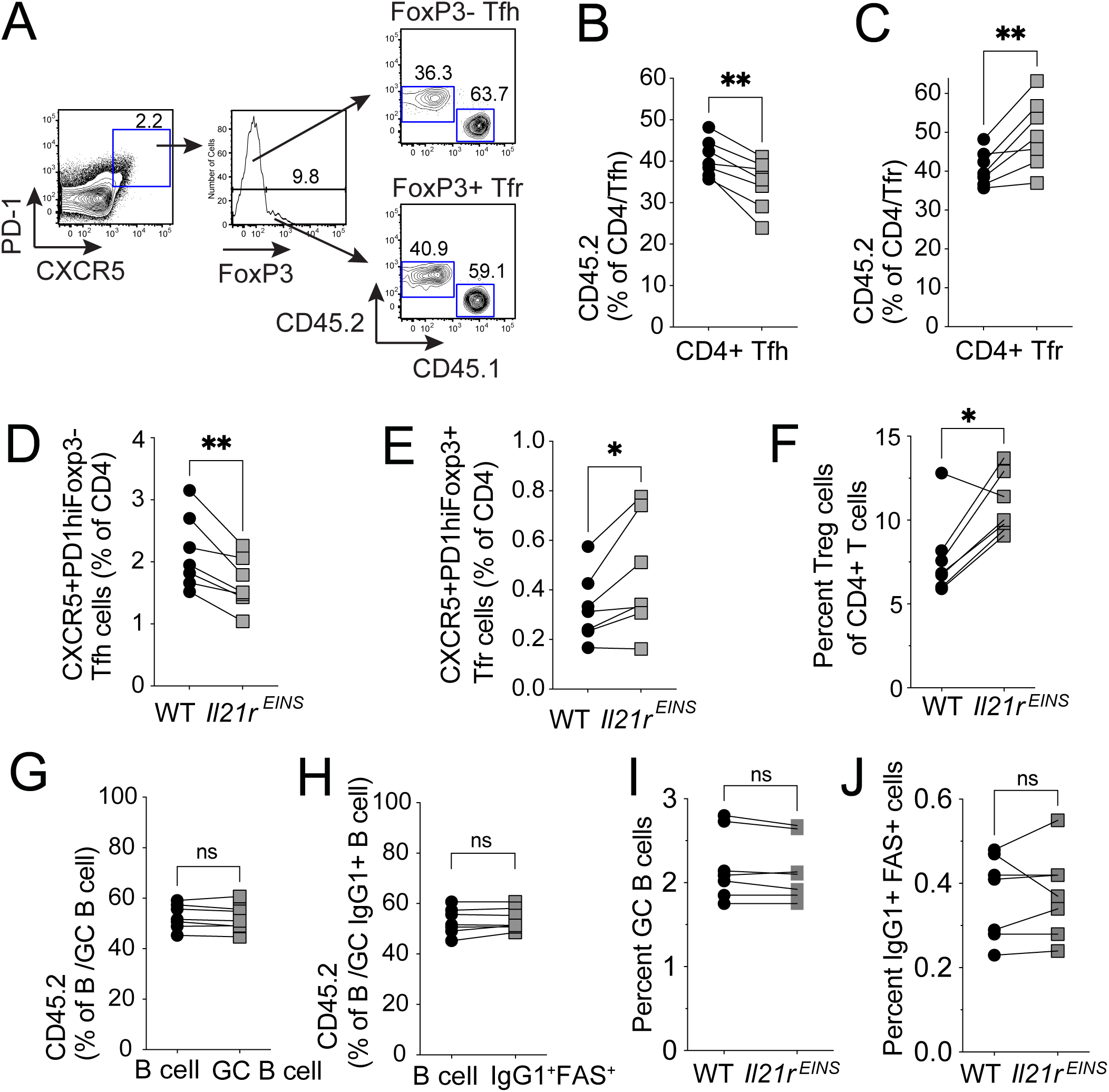
T cell intrinsic effect of *Il21r^EINS^* on the germinal centre response. Mixed BM chimeras were reconstituted with equal ratios of WT CD45.1^+^ BM cells and *Il21r^EINS^*CD45.2^+^ BM cells. 8 weeks after transfer, the mice were immunized with SRBC and analysed 7 days later. Gating strategy employed to differentiate Tfh and Tfr cells from WT CD45.1^+^ and *Il21r^EINS^* CD45.2^+^ donor mice (A). Percentage of CD45.2^+^ donor cells within the total CD4^+^ T cell population alongside the CD45.2^+^ CXCR5^hi^ PD1^hi^ FoxP3^−^ Tfh cell population (B), percentage of CD45.2^+^ donor cells within the total CD4^+^ T cell population alongside CD45.2^+^ CXCR5^hi^ PD1^hi^ FoxP3^+^ Tfr cells (C). Relative percentage of WT CD45.1^+^ and CD45.2^+^ *Il21r^EINS^* cells of Tfh cells (D), relative percentage of WT CD45.1^+^ and CD45.2^+^ *Il21r^EINS^* cells of Tfr cells (E) relative percentage of CD45.1^+^ versus CD45.2^+^ *Il21r^EINS^* Treg cells (F). Percentage of CD45.2^+^ donor cells within the total B cell population alongside the percentage of CD45.2^+^ donor cells within the GC B cell population (G), percentage of CD45.2^+^ donor B cells within the IgG1^+^ FAS^+^ B cell population (H). Relative percentage of WT CD45.1^+^ and *Il21r^EINS^* CD45.2^+^ GC B cells (I), relative percentage of CD45.1^+^ and CD45.2^+^ IgG1^+^ FAS^+^ B cells (J). Data shown as individual mice n=5 from 3 separate experiments with similar results. Statistical significance was assessed by students T test; *\*p < 0.05; **p < 0.01; ***p < 0.001; ****p < 0.0001*.

Parallel analyses of the donor B cell populations after immunization with SRBC showed that within the total B cell population approximately 53.1% are CD45.2^+^ *Il21r^EINS^* (Fig. 4G). In contrast to what was observed for the *Il21r^EINS^* T follicular cell subsets, we did not detect a cell intrinsic effect of the *einseitig* mutation in either GC B cells (Fig. 4G) or IgG1^+^ FAS^+^ B cells (Fig. 4H). Similarly, comparison of B cell subsets derived from *Il21r^EINS^*and WT donors’ side by side showed no difference between *Il21r^EINS^*GC B cells (Fig. 4I) or IgG1^+^ FAS^+^ B cells (Fig. 4J) in comparison with WT GC B cells. Collectively, our findings indicate a cell intrinsic contribution of IL21-dependent STAT1-signaling to the differentiation of T follicular subsets, but not GC B cells.

### Reduced IL-21 dependent STAT1 phosphorylation leads to quantitative and qualitative differences in follicular CD4^+^ T cell populations

Whilst the frequency of Tfh cells were reduced by the *einseitig* mutation, it was unclear whether the expression of Tfh functional molecules was also affected. IL-21R signalling within Tfh cells increases the proportion of Tfh cells that produce IL-21 *in vivo*, thereby establishing a feed-forward loop (*14*). When we assessed IL-21 production in Tfh cells 7 days after SRBC immunization, we observed a decrease in the percentages of IL-21^+^ Tfh cells in *IL21r^EINS^*in comparison to their WT counterparts (Fig. 5A). Furthermore, the levels of IL-21 produced by *Il21r^EINS^* Tfh cells were decreased when compared with WT Tfh cells (Fig. 5B and Fig. 5C).

**Figure 5.**
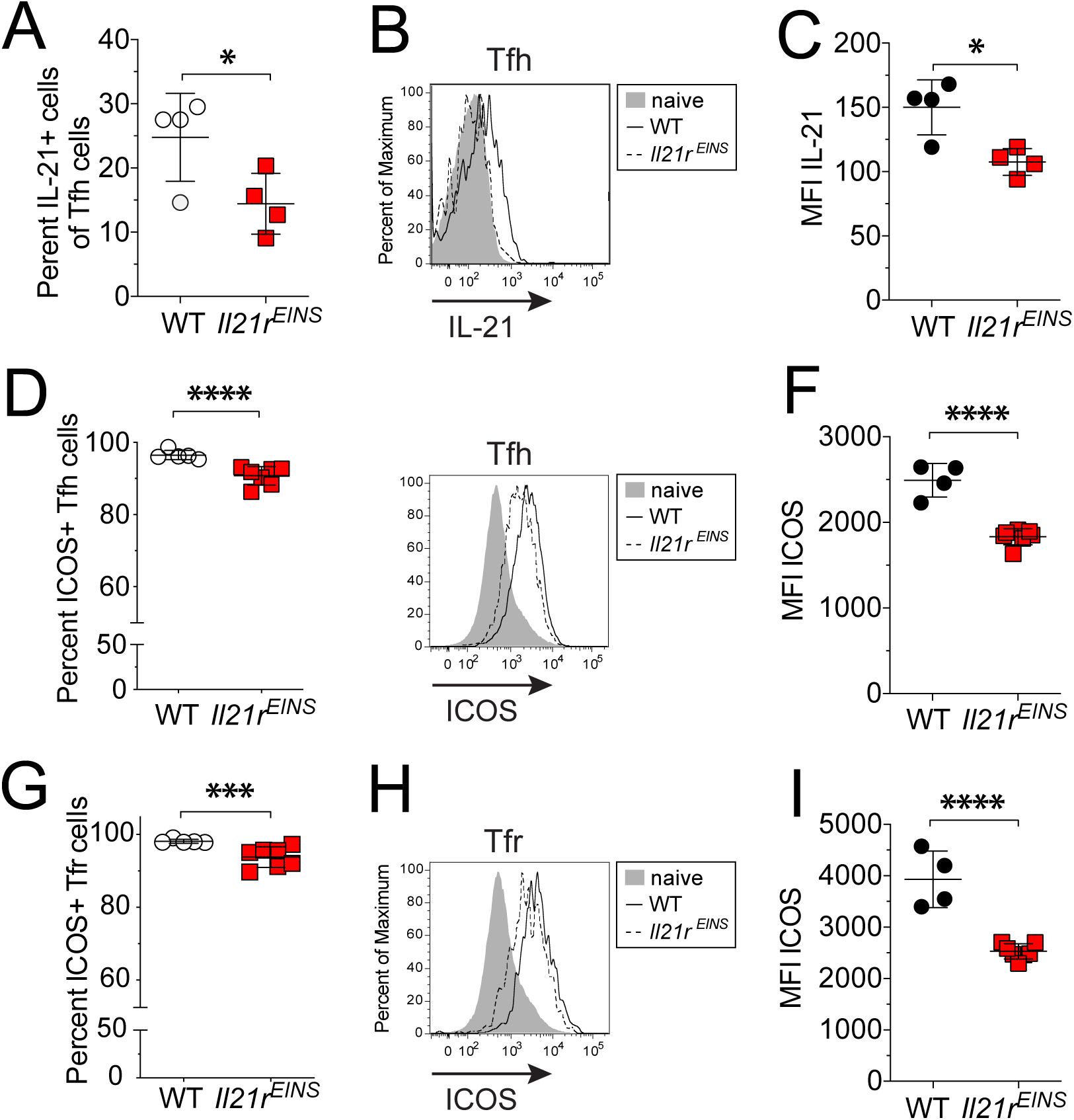
The *Il21r^EINS^* mutation affects the expression of Tfh functional molecules. Staining for ICOS and IL-21 on Tfh and Tfr cells from WT and *Il21r^EINS^* 7 days after SRBC immunization. Percentage of IL-21+ Tfh cells (A), histogram showing IL-21 expression in Tfh cells (B), quantification of IL-21 expression (MFI) in Tfh cells (C). Percentage of ICOS^+^ Tfh cells (D), Histogram showing ICOS expression on Tfh cells (E), levels of ICOS expression on Tfh cells (MFI) (F). Percentage ICOS^+^ Tfr cells (G), histogram showing ICOS expression on Tfr cells (H), levels of ICOS expression on Tfr cells (I). Data are shown from one representative experiment as individual mice and means +/- SD, from 3 experimental replicates with similar results. Statistical significance was assessed by students T test; *\*p < 0.05; **p < 0.01; ***p < 0.001; ****p < 0.0001*.

T follicular cells express the costimulatory molecule inducible T cell co-stimulator (ICOS), which is important for Tfh cell differentiation and function in humans and mice (*38, 39*). The vast majority of WT Tfh cells express ICOS, which were reduced in *Il21r^EINS^* CXCR5^hi^ PD1^hi^ CD4^+^ Tfh cells (Fig. 5D) and, more notably, the intensity of ICOS-expression was lower on *Il21r^EINS^* Tfh cells compared with WT Tfh cells (Fig. 5E and Fig. 5F). The observed decrease in ICOS expression was not limited to the follicular helper population, since the ICOS-expressing fraction of the *Il21r^EINS^* Tfr population was also reduced relative to WT cells (Fig. 5G), as well as the levels of ICOS-expression on *Il21r^EINS^* Tfr cells (Fig. 5H and Fig. 5I). These observations demonstrated that IL-21 induced STAT1-signaling was important for optimal IL-21 production by Tfh cells and for the expression of ICOS on T follicular cell subsets. Therefore, IL-21:IL-21R induced STAT1 activation plays an important role in both the differentiation and function of Tfh cells.

### IL-21 induced activation of STAT1 does not control CD4^+^ T cell expression of IL-6Rα

As discussed earlier, IL-6 is produced following SRBC immunization (*29*) and has been shown to compensate, to varying degrees, for IL-21 during the differentiation of *Il21r^−/−^* Tfh cells (*16, 21*). Both cytokines play a role in Tfh cell differentiation, shown by the severe reduction of Tfh cells in mice made deficient in both IL-21 and IL-6 in response to virus infection (*16*). Therefore, we were intrigued that whilst IL-6 can compensate for Tfh cell differentiation in the absence of IL-21R, it could not compensate in the presence of a mutant IL-21R, potentially due to a niche-filling effect of the mutant IL-21R (*40*). To test this, we first assessed, expression of the alpha chain of the IL-6 receptor (IL-6Rα) on CD4^+^ T cells. Flow cytometric analyses indicated increased expression of IL-6Rα (Fig. 6A) and an increased percentage of IL-6Rα expressing *Il21r^−/−^* CD4^+^ T cells (Fig 6B), compared with both *Il21r^EINS^*and WT CD4^+^ T cells.

**Figure 6.**
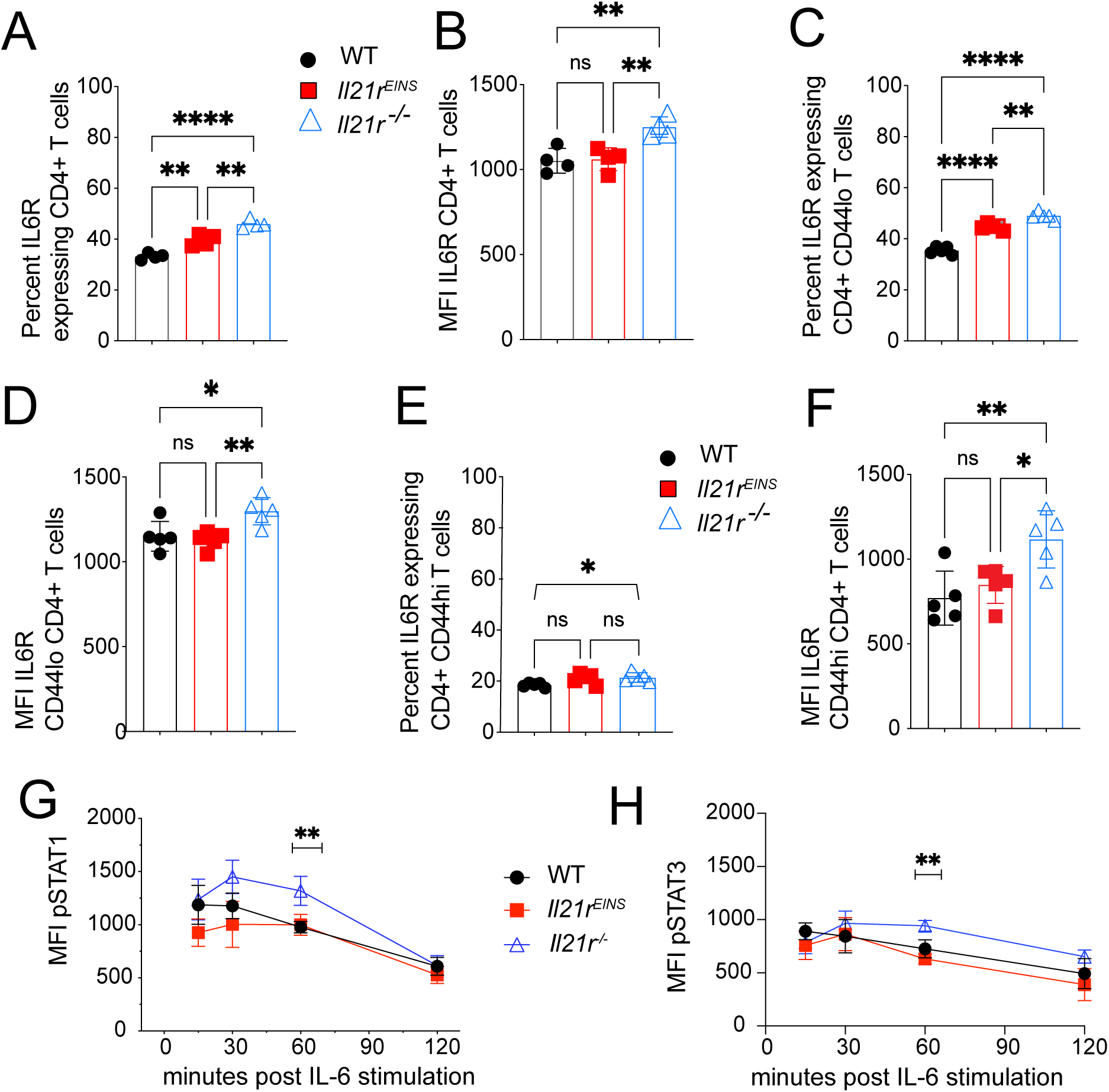
Genetic loss of IL-21:IL-21R signalling, but not IL-21 mediated STAT1 activation, drives IL-6Rα expression and responsiveness to IL-6. IL-6 receptor (IL-6Rα) expression on CD4^+^ T cells from the spleen of WT, *Il21r^−/−^*and *Il21r^EINS^* mice. Percentage of IL-6Rα expressing CD4^+^ T cells (A) and mean fluorescence intensity (MFI) of IL-6Rα on CD4^+^ T cells (B). Percentage (C) and MFI (D) of IL-6Rα expressing naïve CD44^lo^ CD4^+^ T cells. Percentage (E) and MFI (F) of IL-6Rα expressing activated/memory phenotype CD44^hi^ CD4^+^ T cells. Graphs show data for individual mice, means +/- SD from two experiments with similar results. Statistical significance was assessed by one-way ANOVA using Bonferroni’s multiple comparisons test; *\*p < 0.05; **p < 0.01; ***p < 0.001; ****p < 0.0001.* Flow cytometric analyses of the percentage of CD4^+^ T cells from the spleen of WT, *Il21r^−/−^*and *Il21r^EINS^* mice containing phosphorylated STAT1 (G) and phosphorylated STAT3 (H) after stimulation with 40 ng/ml rmIL-6 over a 2-hour time course detected by immunostaining of phosphorylated STAT proteins, flow cytometry and FACS analyses. Statistical significance was assessed by two-way ANOVA with multiple comparisons; *\*p < 0.05; **p < 0.01; ***p < 0.001; ****p < 0.0001*.

Since IL-6Rα is expressed on both CD44^lo^ naïve and CD44^hi^ activated/memory phenotype CD4^+^ T cells, we analysed naïve CD4^+^ T cells within the total CD4^+^ T cell population in the spleen, which demonstrated a greater percentage of IL-6Rα expressing *Il21r^−/−^* CD4^+^ T cells than WT cells or *Il21r^EINS^* cells (Fig. 6C). Whereas *Il21r^EINS^* IL-6Rα^+^ naïve cells were intermediate; with a greater percentage relative to WT cells, but less than *Il21r^−/−^* cells (Fig. 6C). By contrast, the levels of IL-6Rα expression were higher on naïve CD4^+^ T cells that lack IL-21:IL-21R signalling relative to both WT and *Il21r^EINS^*cells (Fig. 6D). In all genotypes, the expression of IL-6Rα was lower on activated/memory phenotype CD4^+^ T cells than naïve as has been shown previously (*41*). However, IL-6Rα expressing *Il21r^−/−^* CD44^hi^ CD4^+^ T cells, in turn, were even further reduced when compared with WT, but not *Il21r^EINS^* CD44^hi^ CD4^+^ T cells (Fig. 6E). Similar to naïve CD4^+^ T cells, the levels of IL-6Rα expression on *Il21r^−/−^* CD44^hi^ cells were markedly higher when compared with both *Il21r^EINS^* and WT CD44^hi^ CD4^+^ cells (Fig. 6F). Taken together, these findings indicate that IL-21 mediated STAT1 activation affected the percentages of IL-6Rα expressing naïve CD4+ T cells but not the level of expression of IL-6Rα on CD4^+^ T cells.

In classic IL-6 signalling, the soluble cytokine binds to its membrane bound IL-6Rα, which then associates with signal-transducing β-receptor (gp130) resulting in the activation of STAT1 and STAT3. To test responsiveness to IL-6, we first assessed Th17 differentiation, which is strongly dependent on signals from IL-6 (*42*), by culturing WT, *Il21r^−/−^* and *Il21r^EINS^*CD4^+^ T cells under Th17 polarizing conditions. On day 3 of culture, we observed similar percentages of *Il21r^EINS^* and WT IL-17^+^ Th17 cells, but reduced differentiation of *Il21r^−/−^* Th17 cells, as has been previously reported (*43, 44*) indicating that the biological function of IL-6 was intact in *Il21r^EINS^* CD4^+^ T cells (Figure S5). Whilst IL-6 was able to support Th17 differentiation in *Il21r^EINS^*CD4^+^ T cells, Th17 differentiation was reduced in *Il21r^−/−^*CD4^+^ T cells despite increased levels of IL-6Rα. These observations suggested that responsiveness to IL-6 may be altered in the absence of normal IL-

21:IL-21R signalling. Analyses of the activation of STAT1 and STAT3 in WT, *Il21r^−/−^* and *Il21r^EINS^* CD4^+^ T cells exposed to low dose rmIL-6 indicated that each genotype responded similarly (Figure S5). We then extended these analyses to test the level of STAT1 and STAT3 activation in WT, *Il21r^−/−^*and *Il21r^EINS^* CD4^+^ T cells over a 2-hour time course. These findings demonstrated that *Il21r^−/−^* CD4^+^ T cells maintained higher levels of phosphorylated STAT1 (Fig. 6G) and STAT3 (Fig. 6H) at 60 minutes following exposure to rmIL-6. By contrast, WT and *Il21r^EINS^* CD4^+^ T cells showed equivalent levels of both phosphorylated STAT1 (Fig. 6G) and STAT3 (Fig. 6H). These findings indicate that the absence of the IL-21R, but not the presence of an IL-21R STAT1 hypomorph, alters the kinetics or maintenance of STAT1 and STAT3 activation over time in response to IL-6.

### IL-6 is unable to compensate for IL-21 in the presence of a IL-21R STAT1 hypomorph

The observation of increased expression of IL-6Rα on *Il21r^−/−^* CD4^+^ T cells and delayed STAT1 and STAT3 activation in response to IL-6 may refect differences in the capacity to ulitize IL-6 in the absence of IL-21:IL-21R signalling. To test whether IL-21 mediated STAT1 activation affected IL-6 mediated compensation, we utilized a previously published method (*24*) to block IL-6 signalling *in vivo* by treating WT, *Il21r^EINS^* and *Il21r^−/−^* mice with neutralizing anti-IL6 monoclonal antibody (mAb). Analyses of untreated mice 7 days after SRBC immunization showed that the percentage of *Il21r^−/−^* CXCR5^hi^ PD1^hi^ T follicular cells expressing IL-6Rα was comparable between genotypes (Fig. 7A), but the level of expression of IL-6Rα was greater on *Il21r^−/−^*Tfh cells compared with WT and *Il21r^EINS^* cells (Fig. 7B and Fig. 7C).

**Figure 7.**
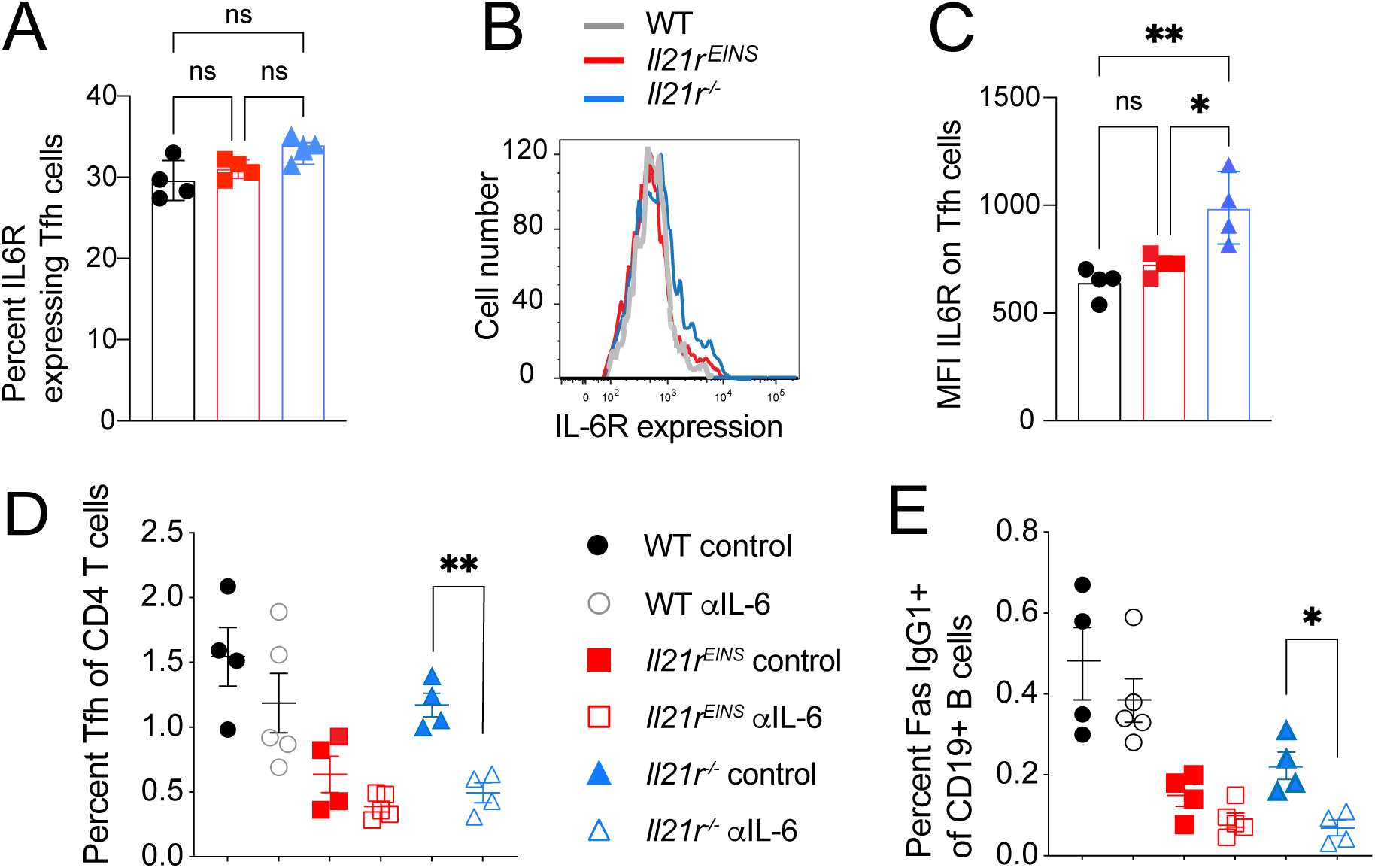
The effect of *Il21r^EINS^* on the contribution of IL-6 to the GC response. Analysis of the germinal centre (GC) response in the spleen in WT, *Il21r^−/−^* and *Il21r^EINS^* mice on day 7 following SRBC immunization. IL-6 receptor (IL-6Rα) expression on CXCR5^hi^ PD1^hi^ FoxP3^−^ T follicular helper (Tfh) cells from the spleen of WT, *Il21r^−/−^* and *Il21r^EINS^* mice showing percentage (A) and mean fluorescence intensity (MFI) (B) and quantitation of MFI of IL-6Rα on Tfh cells (C). IL-6 signalling was blocked *in vivo* by the injection of 0.5mg neutralizing anti-IL-6 mAb or isotype mAb and 0.25mg on day 0, the day of SRBC immunisation, and then day 2, day 4 and day 6. GC populations were analysed in the spleen on day 7. (D) Percentage of Tfh cells of CD4^+^ T cells, (E) percent IgG1^+^FAS^+^ B cells of total B cells. Data shown as individual mice n=5 from 2 experimental repeats with similar results. Statistical significance was assessed by 1-way ANOVA using Bonferroni’s multiple comparisons test; *\*p < 0.05; **p < 0.01; ***p < 0.001; ****p < 0.0001*.

Neutralisation of IL-6 with anti-IL6 mAb did not significantly reduce the FoxP3^−^ Tfh cell population in either WT or *Il21r^EINS^* but did reduce the percentage of *Il21r^−/−^* Tfh cells 7 days after immunization with SRBC (Fig. 7D). By contrast, we could not detect a significant change upon *in vivo* neutralization of IL-6 on the FoxP3^+^ Tfr population in either WT, *Il21r^EINS^* or *Il21r^−/−^*mice (Figure S5). Similarly, we did not observe a significant difference in the percentage of GC B cells in WT, *Il21r^EINS^* and *Il21r−/−* mice (Figure S5). However, IgG1^+^ FAS^+^ B cells were decreased in *Il21r^−/−^* but not *Il21r^EINS^* or WT mice upon *in vivo* blockade of IL-6 (Fig. 7E). Our *in vivo* IL-6 blocking experiments indicated a significantly greater contribution of IL-6 to GC output in *Il21r^−/−^*than *Il21r^EINS^* or WT mice. Taken together these findings demonstrate that, despite the strong ability of IL-6 to activate STAT1, IL-6 is ineffective at fully compensating for the geminal centre response in the presence of an IL-21 receptor mutant that predominantly affects IL-21 activation of STAT1.

## Discussion

The importance of IL-21 induced JAK-STAT signalling for the humoral immune response to T-dependent antigen has been confirmed by a wide body of data (*10–13, 31*). However, the relative role of IL21-dependent STAT1, STAT3 and STAT5 activation on the GC response has been difficult to tell apart. Here, we identify an IL-21R hypomorph and take advantage of its selective defect in STAT1 activation to elucidate the role of IL-21 induced STAT1 activation in the differentiation of GC T and B cells The ENU-induced mutation in *Il21r^EINS^* mice caused a change from a negatively charged Aspartic acid to a neutral, hydrophobic, Valine at the highly conserved amino acid position 375 in the cytoplasmic tail region of the IL-21R receptor protein. Whilst previous studies have demonstrated that the main STAT1 docking tyrosine is Y510 (*8*), the *Il21r^EINS^* mutation may affect the geometry or stability of the IL-21R, leading to diminished STAT1 phosphorylation. Comparison of the pSTAT1 levels of IL-21 stimulated cells from *Il21r^EINS^* and WT mice revealed diminished STAT1 activation by mutant IL-21R in CD4^+^ T cells, whereas STAT3 and STAT5 activation were unaffected. In contrast to the STAT3 dominance observed following interferon-gamma stimulation of *STAT1* deficient cells (*31*), we did not observe a compensatory increase in STAT3 phosphorylation in *Il21r^EINS^* CD4^+^ T cells. Moreover, activation of STAT1 by cytokines acting through different receptors, such as IL-6, remained undisturbed in *Il21r^EINS^* CD4^+^ T cells.

Previous studies have demonstrated an important role for STAT1 and STAT3 activation on GC populations. For instance, STAT3 has a nonredundant role in human Tfh cell differentiation (*20*) and IL-21-dependent STAT3 phosphorylation in human B cells is crucial for the development of a strong high-affinity plasmablast response (*7, 18*). In murine CD4^+^ T cells, STAT3 expression is required for both Tfh cell and GC B cell differentiation and generation of virus-specific antibody responses (*22*). IL-6, in turn, mediates STAT1 activation that is important for the induction of Bcl-6 in early Tfh cell differentiation (*21*). This study has revealed a previously unappreciated role for IL-21 mediated STAT1 activation in the both the differentiation of Tfh cells and the expression of molecules important to Tfh cell function. *Il21r^EINS^*Tfh cells exhibited reduced IL-21 production and ICOS expression was reduced on both *Il21r^EINS^* Tfh and *Il21r^EINS^* Tfr cells. This finding indicates that STAT1 activated by IL-21 is important for optimal Tfh cell differentiation.

Analyses of mixed BM chimeras immunised with SRBC, identified a T cell intrinsic contribution for IL-21 dependent STAT1 activation to both the differentiation of Tfh cells and the inhibition of Tfr cells. However, the T cell intrinsic effect was small, and additional factors may contribute to the reduced differentiation of *Il21r^EINS^*Tfh cells, including the effect of IL-21:IL-21R signalling on non-lymphoid compartments. By contrast, there was no B cell intrinsic role for the mutant IL-21R for the differentiation of GC B cell populations. The absence of a B cell intrinsic effect of the reduced IL-21 mediated STAT1 phosphorylation might be explained by a stronger dependency of B cells on IL-21 mediated STAT3 activation (*7, 18, 45*).

We, and others, have previously observed a T cell intrinsic role for IL-21 signalling in the GC response in mice immunised with OVA delivered in alum (*11, 14*). By contrast, immunisation with the T dependent antigen SRBC, that induces ample IL-6 expression (*29*), did not reveal a significant reduction in *Il21r^−/−^*Tfh cells, as has been previously reported (*12*). The observation that IL-6 can act redundantly, to varying degrees, to support *Il21r^−/−^*Tfh cell differentiation (*16, 24, 46*), delivering an important STAT1 mediated signal (*21*), suggested that IL-6 may contribute to Tfh cell differentiation in the absence of the *Il21r* but not in the presence of a defective receptor. This notion was supported by increased expression of IL-6Rα on *Il21r^−/−^* CD4^+^ T cells compared with WT cells with an intact IL-21R or an *Il21r^EINS^* hypomorph.

IL-6Rα is highly expressed on naïve and central memory T cells (*41, 47*), but decreases as IL-6Rα undergoes cleavage following T cell activation and IL-6 ligation, resulting in the release of soluble IL-6Rα (*41*). In this context, the increased levels of IL-6Rα on *Il21r^−/−^* naïve and memory phenotype CD4^+^ T cells in unmanipulated mice might suggest a reduced level of T cell activation in *Il21r^−/−^* mice. The reason for increased expression of IL-6Rα on cells that lack IL-21:IL-21R signalling is not known, but may reflect IL-21’s role as a costimulatory molecule and that IL-21 and IL-6 each work optimally when both cytokines are present (*28, 48*). Despite clear differences in the level of expression of IL-6Rα on *Il21r^−/−^* CD4^+^ T cells, we did not find a marked difference in the levels of either activated STAT1 or STAT3 in response to stimulation with IL-6 *in vitro*. However, we did observe that *Il21r^−/−^* CD4^+^ T cells had a delayed peak, or more sustained levels, of phosphorylated STAT1 and STAT3 over time and were more affected by neutralization of IL-6 *in vivo*. This indicates that expression of IL-6Rα is affected by IL-21:IL-21R signalling, which may influence utilization of IL-6 in *Il21r^−/−^* CD4^+^ T cells following SRBC immunization.

Identification of the *Il21r^EINS^* mutation has enabled the study of the distinct contribution of IL-21 mediated STAT1 activation in the germinal centre response. Despite the potential input of other STAT1 inducing cytokines during T dependent immunisation, IL-21 induced STAT1 activation could not be entirely compensated for. Functional compensation has been reported in several settings to explain differences in the observed effects of genetic deletion versus protein mutants or gene knockdown (*49–52*). Thus, whilst IL-6 has been shown to compensate for a lack of IL-21 in Tfh cell differentiation (*16, 24*), no such redundancy operated to compensate for a IL-21R STAT1 hypomorph. The remaining function of the IL-21^EINS^ receptor likely reduced the need for compensatory pathways that are activated in complete protein knockouts. Future experiments that restore IL-21 induced STAT1 signaling in *Il21r^EINS^* CD4^+^ T cells would provide definitive evidence linking the *Il21r^EINS^* mutation to impaired STAT1 activation in germinal center responses.

## Materials and Methods

### Mice

C57/BL6 (B6), Ly5.1 congenic mice (B6 background), as well as *RAG1^−/−^* mice bred to B6 N10 were purchased from the Animal Research Centre in Perth, Australia. *Il21r^−/−^* mice were obtained from Dr Warren Leonard (NIH) via Dr Mark Smyth (Melbourne) at B6 N6 and backcrossed to N12 for experimental use. *Il21r^EINS^* mice were purchased from the Australian Phenomics facility ANU (Prof Christopher Goodnow). The Garvan Institute and St. Vincent’s Hospital Animal Ethics Committee granted permission to perform animal experiments. Animals were housed under specific pathogen-free conditions and handled in accordance with the Australian code of practice for the care and use of animals for scientific purposes. Age matched littermate mice used for experimental purposes were between 8-14 weeks of age.

### Immunisations and IL-6 neutralisation

Mice were immunized (i.v.) with 2×10^8^ sheep red blood cells (IMVS, Australia) and spleens were analysed at timepoints shown. For *in vivo* neutralization of IL-6: Mice initially received 0.5 mg neutralizing anti-IL-6 (clone MP5-20F3, rat IgG1; BioXCell) or isotype mAb (anti-rat IgG1; BioXCell), and subsequently given 0.25 mg mAb (i.v.) every other day following the initial dose through the end of the experiment (*24*).

### Bone Marrow Chimeras

Cohorts of *RAG1^−/−^* mice were lethally irradiated by a ^137^Cs source (single dose of 0.6Gy) and reconstituted the following day by i.v. injection with BM cells isolated from femurs and tibiae obtained by flushing bone with lymphocyte isolation media in sterile conditions. After irradiation, mice were maintained on antibiotic water containing cotrimoxazole (Roche) for 7 days. Irradiated *RAG1^−/−^* mice were reconstituted with 1×10^7^ BM cells from Ly5.1 congenic mice (B6 background) expressing the congenic marker CD45.1 to allow detection, combined with equal proportions BM from CD45.2^+^ WT or CD45.2^+^ *Il21r^EINS^* mice. Since no mature lymphocytes develop in *RAG1^−/−^*mice, no radio-resistant donor T cells were present in our analysis. Our reconstitution protocol typically resulted in an approximate 35-65:65-35 ratio of WT:*Il21r^EINS^*lymphocytes. Mice were used for immunization protocols as described above at eight weeks post-reconstitution.

### Immunofluorescence and FACS analyses

For flow cytometric analysis of mouse lymphocytes, red blood cells (RBC) were removed by hypotonic lysis. 50μl of a single cell suspension at 2×10^7^ cells/ml from spleen and lymph nodes were stained in FACS buffer containing pre-diluted antibodies in 96 well V-bottomed microtitre plates (Nunc, Roskilde, Denmark). Antibodies against surface molecules purchased from BD Bioscience were anti-B220-APC & APC-Cy7 (RA3-6B2, dilution 1/300), anti-CD4-PE-Cy7 & A488 (RM4-5, dilution 1/300), anti-CD21-FITC (7G6, dilution 1/200), anti-CD23-PE (B3B4, dilution 1/200), anti-CD25-APC & BV605 (PC61, dilution 1/100), anti-CD45.1-PE-Cy7 (A20, dilution 1/200), anti-CD45.2-APC-Cy7 (104, dilution 1/200), anti-CXCR5-biotin (2G8, dilution 1/100), anti-GL7-PE (GL7, dilution 1/300), anti-FAS-FITC (Jo2, dilution 1/200), anti-IgG1-biotin (A85-1, dilution 1/100) and anti-Ly6G-PB (1A8, 1/300). Antibodies purchased from eBioscience were anti-CD8-APC (53-6.7, dilution 1/200), anti-CD11b-FITC (M1/70, dilution 1/200), anti-CD11c-APC (N418, dilution 1/200), anti-CD19-eF450 (1D3, dilution 1/200), anti-CD38-PE-Cy5.5 (90, dilution 1/200), anti-CD44-APC-A780 (IM7, dilution 1/200), anti-ICOS-PE (C398.4A, dilution 1/300),anti-NK1.1-PE (PK136, dilution 1/200), anti-PD-1-FITC (RMP1-30, dilution 1/100), Streptavidin-PerCP-Cy5.5 & APC (dilution 1/300), anti-TCRβ-APC, FITC & PerCP-Cy5.5 (H57-597, dilution 1/300). Intracellular immunostaining for FoxP3 carried out using the anti-Foxp3-eF450 (FJK-16s, dilution 1/100) Antibodies against intracellular proteins were purchased from eBioscience using the BD intracellular staining kit (Cat#554714) according to the manufacturer’s instructions. Intracellular phosphorylated STAT proteins were detected with anti-STAT1 (PY701)-A647 (4a, dilution 1/5), anti-STAT3 (PY705)-PE (4/P-STAT3, dilution 1/5) and anti-P-STAT5-PerCP-Cy5.5 (47, dilution 1/5) mAbs using the BD Phosphlow buffer III (Cat#558050) according to manufacturer’s instructions. Cytokines were detected either directly ex-vivo or after 4 hours stimulation at 37°C in cell culture media with Phorbol 12-myristate 13 acetate (PMA 500ng/ml, BIOMOL USA), ionomycin (500ng/ml, Invitrogen) and GolgiStop (1:1000, BD Biosciences, CA). Intracellular IL-21 was detected using IL-21rFc (manufactured in house), conjugated to A647 using the Molecular Probes A647 Antibody Labeling Kit (Thermo Fisher) according to the manufacturer’s instructions. Intracellular IL-17 was detected using anti-IL-17a-APC-Cy (TC11-18H10, dilution 1/100) purchased from BD Biosciences and staining for intracellular cytokines was carried out using the BD intracellular staining kit (Cat#554714) according to the manufacturer’s instructions. Cells were acquired using either a FACS Canto II or a Fortessa cytometer (BD Biosciences, CA) and analysed using Flowjo (Treestar, CA).

### Immunohistochemistry

5μm frozen OCT (Tissue Tek, Australia) spleen sections were fixed in ice-cold acetone for 8 minutes, washed and incubated with blocking reagent (Avidin/biotin (Dako)) at room temperature. Primary biotin or fluorochrome-conjugated antibodies were incubated at RT overnight. Abs against surface molecules purchased from BD Biosciences were anti-CD4-PE (RM4-5, dilution 1/100) and anti-IgG1-biotin (A85-1, dilution 1/100). Streptavidin-APC (dilution 1/200) was purchased from eBioscience and IgD was detected using anti-IgD-A488 (11-26c2a, dilution 1/100) Ab purchased from Biolegend. Sections were analysed using a Leica DM I6000 SP8 confocal microscope (Leica Microsystems, Wetzlar, Germany). The images were processed using the Leica acquisition and analysis software ImageJ (Freeware NIH Bethesda, USA) or Adobe Photoshop, version 7 (San José, CA).

### SDS Page and Immunoblotting

Splenocytes were stimulated at a concentration of 3×10^6^ cells/ml with rmIL-21 (Peprotech, NJ) at 80ng/ml for or incubated in media 15 minutes. Cells were washed and lysed in RIPA buffer containing protease and phosphatase inhibitors (SIGMA, CA) and run by SDS polyacrylamide-gel electrophoresis using a 4-12% gradient gel and nitrocellulose membranes (Invitrogen). Membranes were probed with anti-STAT1 (42H3, Cell Signalling, USA) or anti-STAT3 (F-2, Santa Cruz, USA) and anti-β-actin (AC-74, Sigma Aldrich, USA). Antibody binding was detected using goat anti-rabbit IgG-HRP (cat# SC-2004, Santa Cruz, USA), goat anti-mouse IgG-HRP (cat # 31439, Thermo Fisher, USA), or rabbit anti-goat IgG-HRP (cat# 31433, Thermo Fisher, USA) and developed using ECL (Perkin-Elmer, MA). After the first round of detection membranes were stripped using stripping buffer (0.015% glycine, 0.001% SDS, 0.01% Tween20 in water, pH=2.2) for 30min at 50°C and after two washes with PBS-T re-probed with anti phospho-STAT1 (D4A7, Cell Signalling, USA) or anti phospho-STAT3 (3E2, Cell Signalling, USA).

### Th17 cell conversion in vitro

CD4^+^ T cells from the spleens were isolated by positive selection using magnetic cell sorting by staining CD4^+^ T cells in MACS buffer with anti-CD4 magnetic beads according to the manufacturers instructions and collected using LS columns (both form Milentenyi Biotec, Auburn, USA) giving a cell purity of 90-95%. In brief lymphocytes were resuspended at 2×10^8^ cells/ml and incubated with 10µl anti-CD4 magnetic beads (Milentenyi Biotec, Auburn USA) per 100µl of labelled splenocytes for 20 minutes at 4°C. These cells were washed twice in MACS buffer and CD4^+^ T cells were isolated using LS columns. To induce Th17 cell conversion, CD4^+^ T cells were activated in the presence of 2µg/ml plate-bound anti-CD3 mAb (clone 145-2C11, BD Pharmingen), with soluble 5µg/ml anti-CD28 (clone 37.51, eBioscience), as well as rmIL-6 (50ng/ml, Peprotech,

USA), rmIL-23 (5ng/ml, Peprotech, USA), recombinant human TGFβ1 (1ng/ml, Peprotech, USA), anti-IL-4 mAb (10µg/ml, BD Bioscience) and anti-IFNγ mAb (10µg/ml, BD Bioscience) for 3 days. On day 3, cells were washed in PBS and subsequently stimulated for 4 hours with 500ng/ml PMA (BIOMOL, USA) and 500ng/ml ionomycin (Invitrogen, USA) in the presence of Golgi Stop (1:1000, BD Biosciences, CA). Conversion efficiency was assessed by flow cytometry on day 3 after surface staining for CD4 followed by intracellular staining for IL-17a.

### *In vitro* stimulation

For intracellular pSTAT staining via flow cytometry: 5×10^7^ splenocytes were stimulated at a concentration of 3×10^6^ cells/ml with 80ng/ml rmIL-21 (Peprotech, NJ), rmIL-6 (BD Biosciences, CA) or rmIL-2 (R&D Systems, MN), or at the concentrations indicated, for 15 minutes, or across a 2 hour time course as indicated, before intracellular immunofluorescence staining, flow cytometry and FACS analysis.

## Data analysis and Statistics

Data were analysed using Prism software (Graphpad software, CA) to calculate unpaired, two-way Student’s T test, with an F test to compare variances. Analysis of more than two groups was performed using one-way ANOVA followed by Bonferroni’s test to compare groups or two-way ANOVA as indicated.

## Acknowledgments

This project was supported by project grant funding from NHMRC APP1148051 and Senior Fellowship in the Marie Skłodowska-Curie FCFP at the Freiburg Institute for Advanced Studies (FRIAS), The University of Freiburg, Germany. Our thanks to Professor Bodo Grimbacher Institute for Immunodeficiency, Center for Chronic Immunodeficiency (CCI), Medical Center, Faculty of Medicine, University of Freiburg, Freiburg, Germany.

## Author contributions

Conceptualization: CK, CJ

Methodology: CJ, JW, CK, MB

Visualization: CK, HW, SO

Writing: CK, CJ, HW, SO

Authors declare that they have no competing interests.

## Data and materials availability

All data are available in the main text or the supplementary materials.

## Notes

### Competing Interest Statement

The authors have declared no competing interest.

### Summary of Updates

Thus version of the manuscript has been revised to take reviewer comments into account.

