## Supplementary Figures 1-5 for "An IL-21R STAT hypomorph circumvents functional redundancy in germinal center responses"

**This PDF file includes:**

Figs. S1 to S5

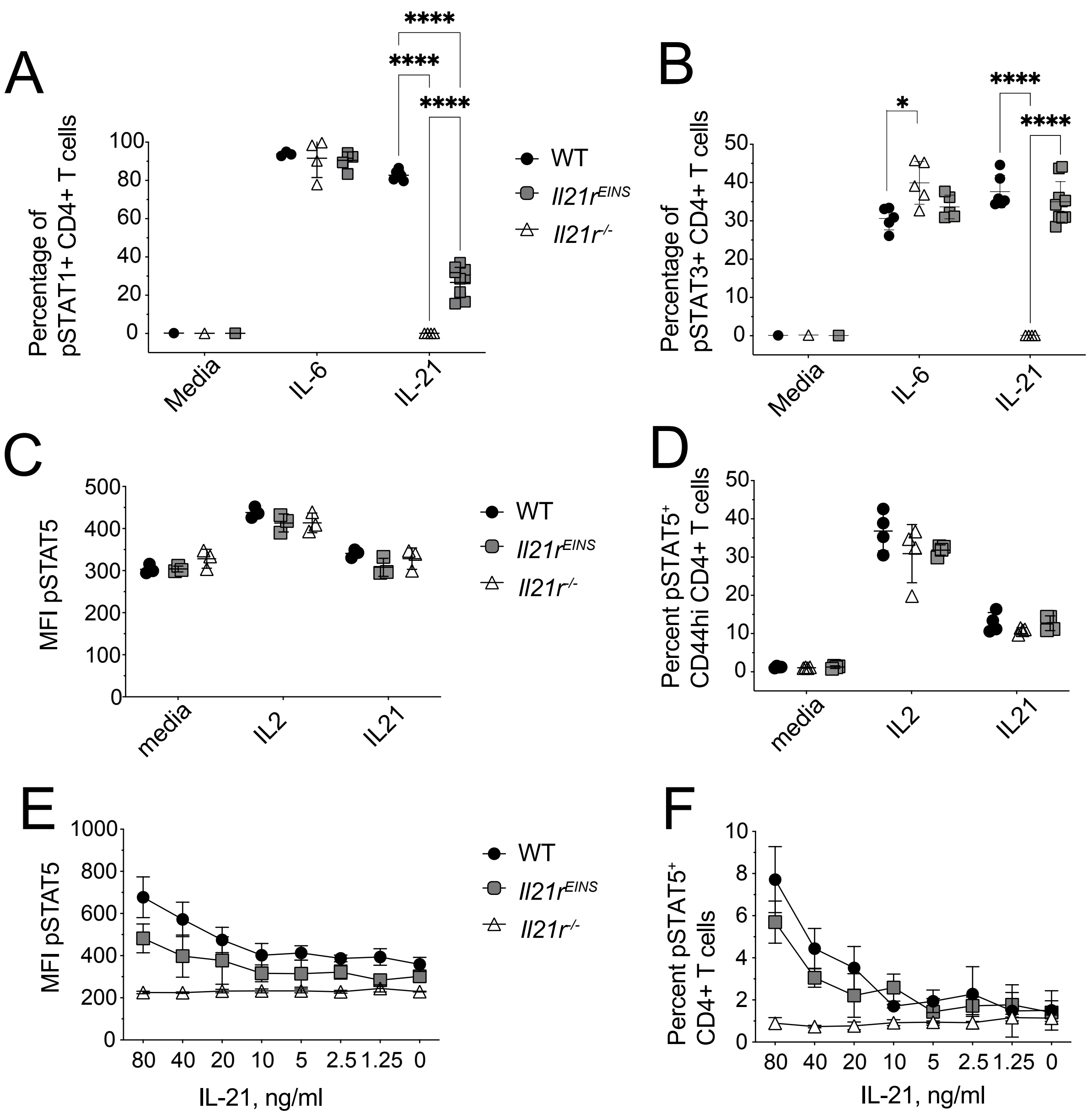


**Supporting Figure 1. Further characterization of the Il21rEINS signaling defect**
Splenocytes from naïve WT, *Il21rEINS* or *Il21r-/-* mice were stimulated with with 100ng/ml rmIL-21, rmIL-6 or rmIL-2 for 15min. Cells were fixed and stained for intracellular phosphorylated (p)STAT3 (A) and pSTAT1 (B) using the BD Phosphlow system and pSTAT expression was analyzed in CD4+ T cells via flow cytometry. Data are shown is representative of 3 similar experiments. Splenocyte derived total CD4+ T cells or CD44hi CD4+ T cells from WT, *Il21rEINS* and *Il21r-/-* mice were stimulated with rmIL-2, rmIL-21 or incubated in media for unstimulated controls for 15min. Cells were fixed and stained for intracellular pSTAT5 using the BD Phosphlow system. Mean fluorescence intensity (MFI) of pSTAT5 in CD4+ T cells (C) and percentage pSTAT5+ CD44high CD4+ T cells (D) were assessed. CD4+ T cells from the spleen of WT, *Il21rEINS* or *Il21r-/-* mice were stimulated with decreasing concentrations of rmIL-21, expression levels of pSTAT5 (E) and percent CD4+ T cells contaiing pSTAT5 (F). Data are shown as mean +/- SD, n= 4-5 mice with 3 experimental replicates. Statistical analyses was performed by 2-way ANOVA.

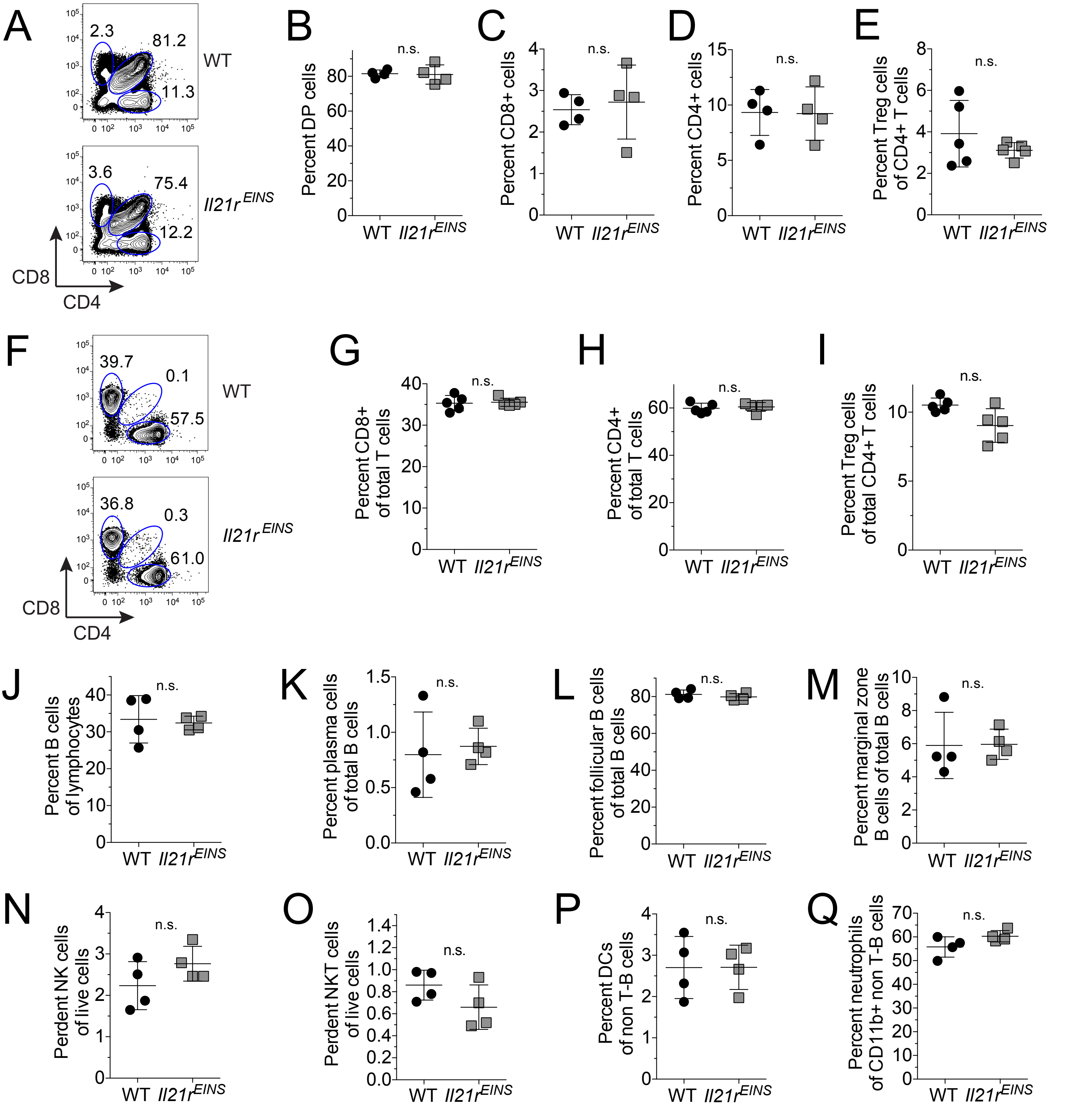


**Supporting Figure 2. Normal lymphocyte development in naïve Il21rEINS mice**

Analysis of thymocytes and splenocytes from naïve WT and Il21rEINS mice, representative dot plot showing the gating of CD4+, CD8+ and CD4+ CD8+ double positive (DP) thymocytes (A), as well as percentages of CD4+ CD8+ DP thymocytes (B), CD8+ single positive thymocytes (C), CD4+ single positive thymocytes (D) and FoxP3+ CD4+ T cells (E) in the thymus. Representative dot plot showing the gating of CD4+, CD8+ T cells in the spleen (F), as well as percentages of CD8+ T cells (G), CD4+ T cells (H) and FoxP3+ CD4+ T cells (I) in the spleen of naïve mice. Also shown are percentages of B cells (J), CD138+ plasma cells (K), CD21med CD23+ follicular B cells (L), CD21+ CD23- marginal zone B cells (M). NK1.1+ NK cells (N), NK1.1+ TCRb+ NKT cells (O), CD11c+ CD11blow TCRβ- B220- dendritic cells (DC) (P) and Ly6G+ CD11c- CD11b+ TCRβ- B220- neutrophils (Q). Values shown from individual mice, including means +/- SD from 3 replicate experiments where n=4-5 per group. Statistical analses perfromed by students T test.

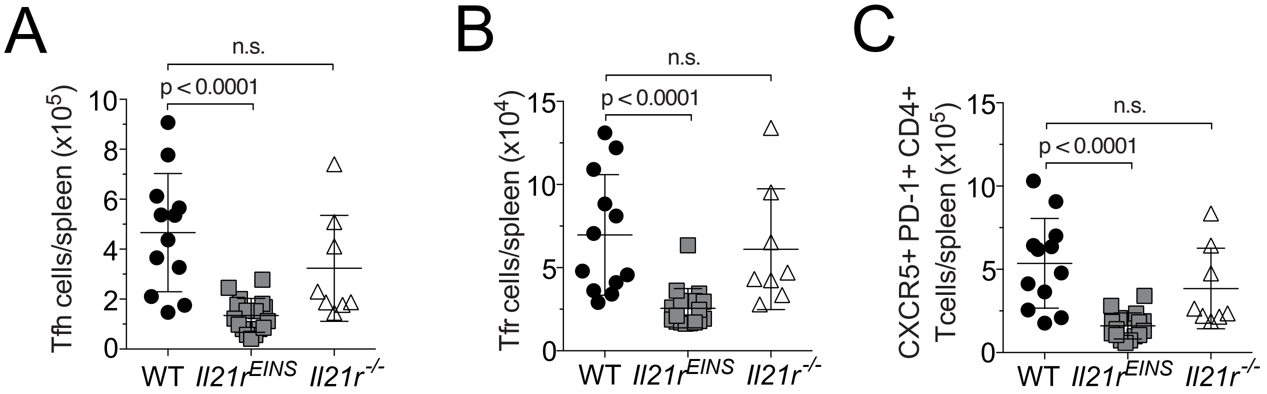


**Supporting Figure 3. Decreased numbers of follicular T cell populations in *Il21r*$^{EINS}$ mice**
*IL21r*$^{EINS}$ mice were immunized wit 2x10^8 SRBCs and lymphocyte populations were analysed 7 days later at the height of the immune response via FACS. Numbers of CXCR5hi PD1hiT FoxP3- follicular helper (Tfh) cells (A), CXCR5hi PD1hiT FoxP3+ T follicular regulatory (Tfr) cells (B), as well as the overall CXCR5+ PD-1+ follicular CD4+ T cell population (C) of WT, *Il21r*$^{EINS}$ and *IL21r-/-* mice are shown. Values shown are from individual mice, including means +/- SD from 3 pooled experiments where n=4-5 per group.

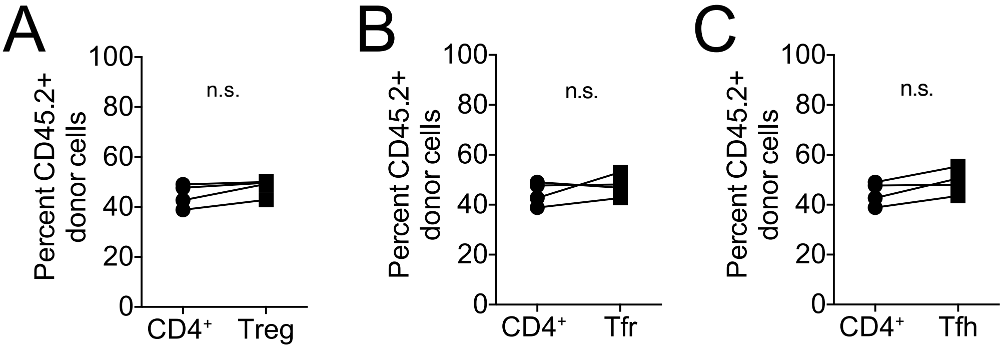


**Supporting Figure 4. CD45.1 WT:CD45.2 WT control mixed bone marrow chimeras**
Control mixed bone marrow (BM) chimeras were reconstituted with equal ratios of C57BL/6 (WT) $CD45.2^{+}$ BM cells and WT $CD45.1^{+}$ BM cells. Eight weeks after transfer, chimeras were immunized with SRBC and analyzed 7 days later. Percentage of $CD45.2^{+}$ donor cells within the total $CD4^{+}$ T cell population shown against: (A) The percentage of $CD45.2^{+}$ donor cells within the Treg population. (B) The percentage $CD45.2^{+}$ within the $CXCR5^{hi}$ $PD1^{hi}$ $FoxP3^{+}$ T follicular regulatory (Tfr) population and (C) the percentage $CD45.2^{+}$ within the $CXCR5^{hi}$ $PD1^{hi}$ $FoxP3^{-}$ T follicular helper (Tfh cell) population. Data shown as individual mice n=5.

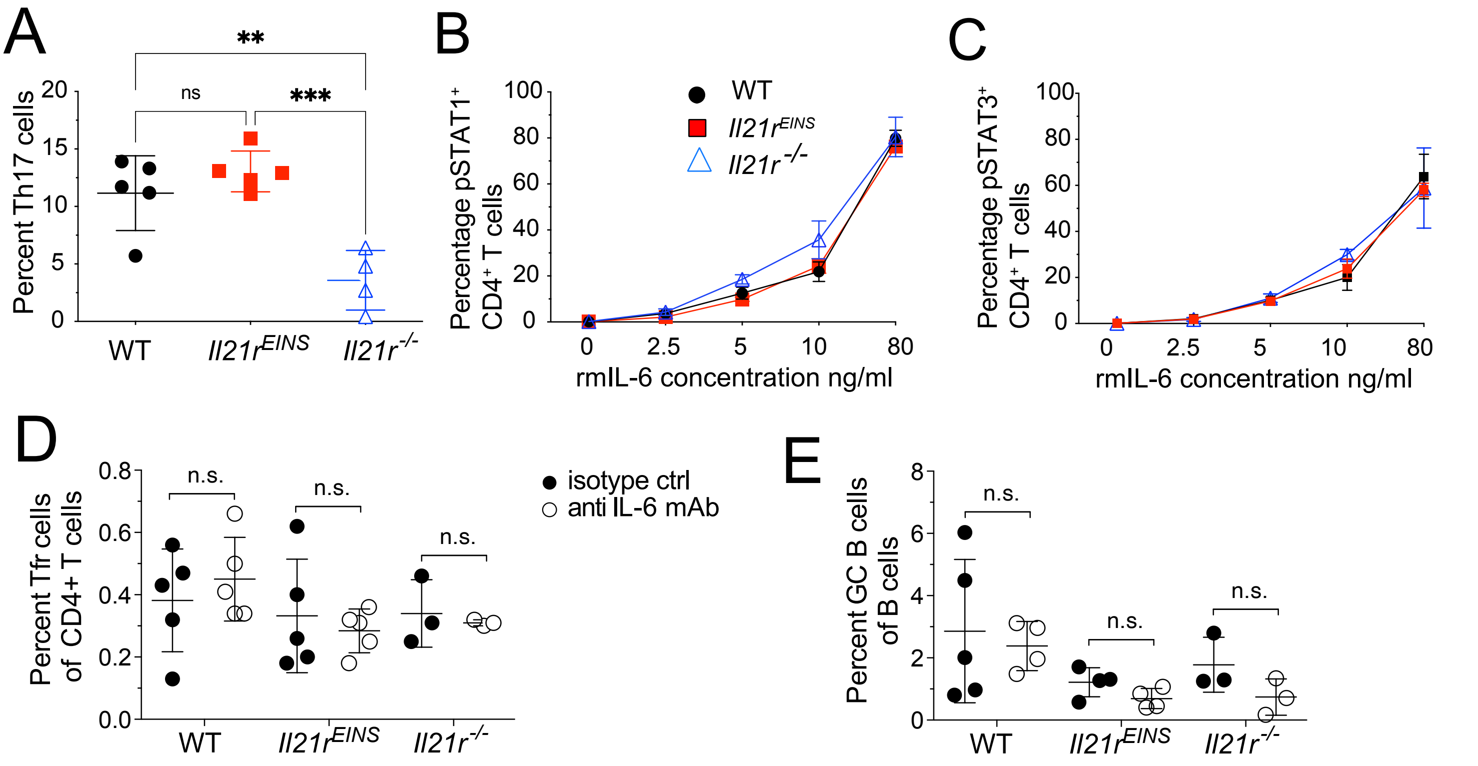


**Supporting Figure 5. IL-6 is still able to carry out its function in IL21rEINS CD4+ T cells**
(A) Th17 cell conversion efficiency was compared using splenic CD4+ T cells form WT and $IL21r^{EINS}$ mice assessed on day 3 of stimulation after 4 hours of restimulation with PMA and ionomycin.IL-17 containing CD4+ T cells were assessed by intracellular immunostaining and flow cytometry. IL-6 signalling was blocked *in vivo* by the injection of 0.5mg neutralizing anti-IL-6 mAb or isotype mAb on day 0 followed by 0.25mg on day 2, day 4 and day 6. Splenocytes from WT, $Il21r^{-/-}$ and $Il21r^{EINS}$ mice were stimulated with rmIL-6 for 15 minutes at the concentrations shown and phosphorylated STAT1 ((B) and STAT3 (C) detected in CD4+ T cells by intracellualr immunostaining and flow cytometry. Mice were immunized with SRBC on day 0 and analyzed via FACS on day 7. Shown here are the percentages of (D) Tfh cells within the CD4+ T cells population and (E) the GC B cell population as a percentage of total B cells. Data are shown as individual mice with n=5 per group from 2 separate experiments with similar results. Statistical significance was assessed by (A-C) one-way ANOVA using Bonferroni's multiple comparisons test and students T test (D and E); *p < 0.05; **p < 0.01; ***p < 0.001; ****p < 0.0001.
